# Bioconvergence of sound-guided and supramolecular assembly strategies to create peptide-protein composite hydrogels with predictable shape-to-function features

**DOI:** 10.1101/2025.06.26.661668

**Authors:** Cosimo Ligorio, Alessandro Cianciosi, Riccardo Tognato, Micaela Natta, Romedi Parolini, Sena Ardicli, Huseyn Babayev, Ieva Sapjanskaite, Samantha L. Kilgour, Richard Homer, Zhiyu Zhou, Andrea Malandrino, Eleni Priglinger, Cezmi Akdis, Martin J. Stoddart, Alvaro Mata, Tiziano Serra

**Affiliations:** Biodiscovery Institute, University of Nottingham, Nottingham, UK; School of Pharmacy, University of Nottingham, Nottingham, UK; Department of Chemical and Environmental Engineering, University of Nottingham, Nottingham, UK; NIHR Nottingham Biomedical Research Centre, Nottingham, NG7 2UH UK; AO Research Institute Davos, Switzerland; Swiss Institute of Allergy and Asthma Research (SIAF), University of Zurich, Davos, Switzerland; Department of Genetics, Faculty of Veterinary Medicine, Bursa Uludag University, Bursa, Turkey; School of Chemistry, University of Nottingham, Nottingham, UK; Wolfson Building, University of Nottingham, Nottingham, UK; Innovation Platform of Regeneration and Repair of Spinal Cord and Nerve Injury, Department of Orthopaedic Surgery, The Seventh Affiliated Hospital, Sun Yat-sen University, Shenzhen, China; Guangdong Provincial Key Laboratory of Orthopaedics and Traumatology, Orthopaedic Research Institute/Department of Spinal Surgery, The First Affiliated Hospital of Sun Yat-sen University, Guangzhou, China; Johannes Kepler University Linz, Kepler University Hospital, Department of Orthopaedics and Traumatology, Linz, Austria; Complex Tissue Regeneration Department, MERLN Institute for Technology-Inspired Regenerative Medicine, Maastricht University, Maastricht, Netherlands

## Abstract

Purely protein-based hydrogels are widely used in tissue engineering for their biomimicry and biocompatibility, yet remain challenging to tailor with precision and predictability at biological and mechanical levels. To overcome this, synthetic self-assembling peptide amphiphiles (PAs) offer opportunities for supramolecular customization, both as single-phase materials and co-assembled with proteins to create hybrid nanocomposites with emerging functionalities. Similarly, contactless, sound-guided bioassembly techniques using liquid-phase hydrogel precursors are emerging as strategic tools for obtaining structured and functional hydrogels. Leveraging these advances, here a fast, contactless, ‘one-pot’ bioassembly strategy merging supramolecular PA self-assembly with sound-guided patterning to fabricate hybrid peptide-protein hydrogels with programmable shape-to-function features is presented. Using fibrin as proof-of-concept, material performance is biologically enhanced by incorporating growth factor-binding PAs, while inorganic microparticles are embedded and spatially organized *via* acoustic fields to tune mechanical properties. This strategy allows predictable tuning of composite stiffness and architecture by adjusting sound wave frequency, with acoustic fields guiding material organization from micro-to-macroscale. Composite hydrogels result highly permissive to cell infiltration *in vitro* and versatile platform to tune immune cell-material interactions. This modular biofabrication platform integrating supramolecular and sound-guided processes can be generalized to other building blocks opening unique opportunities for scalable, tunable, and hierarchically-organized biomaterials.

## Introduction

Purely protein-based biomaterials, such as collagen and fibrin hydrogels, have historically represented a preferred choice for cell culture and tissue engineering, due to their excellent biomimicry and biocompatibility properties. Despite their widespread use, the natural origin of pure proteins makes them a challenging material to be customized with precision at both chemical and mechanical levels. For instance, proteins are known to present multiple bioactive sites, either in full-display or buried in cryptic sites, but the presentation of some epitopes over others remains a challenge due to the inherent structural complexity of proteins^[1]^. Similarly, the use of proteins to rationally-design materials with predictable properties remains difficult due to batch variability, hierarchical assembly, and control over protein structure and protein-protein inter-molecular interactions^[2]^. In addition, other disadvantages include rapid degradation *in vivo* and risks of immunogenicity^[3]^, which further highlight the need for hybridization and creation of multicomponent protein-based biomaterials to achieve tunable functionality and specific biological performance.

Nature has evolved fibrin to serve as the main protein to stop bleeding after trauma and to promote regeneration. Indeed, following rupture of blood vessels, platelets interacting with exposed collagen initiate the coagulation cascade, which ends up with the polymerization of soluble fibrinogen into insoluble fibrin clots. These fibrillary-assembled meshes containing blood cells and key endogenous signals are essential for creating a matrix that stops bleeding, regulates cell infiltration, and ultimately promotes wound healing^[4]^. From a biomaterial perspective, fibrin represents an exceptional protein-based material for wound repair, being bioresorbable, hemostatic, and immunomodulatory. Hence, it is not surprising that fibrin-based materials, such as Evicel^®^, Tisseel^®^, Artiss^®^, and VeraSeal^®^ have been developed as alternatives to metallic staples and polymeric sutures, leading to clinically-approved regenerative products^[5]^.

Despite their widespread use, fibrin-based materials suffer from three major limitations. First, in terms of bioactivity, fibrin strongly relies on interaction partners, such as heparin, to enhance their ability to sequester and release relevant growth factors over time^[6]^. Indeed, in the absence of affinity binders, fibrin gels are often accompanied by fast and uncontrolled drug release, hindering any attempt of precise dosing and delivery^[4]^. Second, fibrin is a soft material characterized by a low mechanical strength (<5 kPa), which is difficult to optimize with precision, as the final mechanical properties strongly depend on intertwining factors^[7]^, such as fibrinogen and thrombin concentrations^[8]^, shear forces^[9]^ and buffers used^[10]^. Finally, fibrin is highly prone to physical contraction and fast degradation over time, making it practically unsuitable for *in vivo* long-term implantation^[11]^. Poor mechanics, absence of binding motifs and rapid degradation highlight the need to effective modification strategies to enhance fibrin performance.

Cross-linking agents or complexes with polymeric and inorganic compounds are often employed to overcome the limitations of fibrin, but with limited success. Among cross-linkers, Factor XIIIa, naturally involved in the coagulation cascade, is currently used to create stiffer fibrin matrices^[12]^, while genipin has been used broadly to cross-link amine-displaying molecules, including fibrin scaffolds^[13,14]^. However, genipin allows tunable cross-linking of fibrin gels but requires careful control of concentration and reaction time to avoid cytotoxicity ^[15]^. Similarly, as Factor XIIIa is enzymatic, it offers less control over cross-linking due to its dependence on substrate availability and reaction kinetics^[16]^. Mechanically, fibrin has been used in composite formulations involving chitosan, silica, acrylates, polyurethanes, and silk. However, despite these efforts, mixtures of fibrin with other components suffer from visible phase separation and heterogeneous gelation, leading to the formation of partially phase-separated continuous networks with poor control over the final mechanical properties^[17]^. Considering these limitations, there is a need for hybrid fibrin materials with precise physico-chemical control.

Enhanced fibrin-based composites with precise properties require fine control over the functionalization and hierarchical organization of fibrinogen with additive molecules. In this direction, at the supramolecular level, self-assembling peptides such as peptide amphiphiles (PAs) can be used to create hybrid peptide-protein materials. By tuning PA structure, PAs have been shown to co-assemble^[18]^ with different types of macromolecules^[19]^ into nanofibers that combine the functionality of both the macromolecules and the peptides.^[20]^ For example, PAs can co-assemble with hyaluronan^[21]^, collagen^[22]^, resilin^[23]^, and elastin-like recombinamers^[24]^ to create peptide-protein structures such as membranes^[21]^, particles with different shapes^[22]^, tubular structures^[25]^ and hydrogels^[26]^ with precise control over their final morphology and mechanical properties^[25]^. Similarly, PAs have been exploited with endogenous ions^[27]^, signaling cytokines and biofluids^[26,28,29]^, to selectively bind, concentrate and release growth factors in a controlled fashion. For these reasons, PAs represent an attractive class of materials to co-assemble with fibrin into supramolecularly-defined peptide-fibrin materials.

At the micro- and macroscale, contactless biofabrication techniques handling low viscosity, liquid precursors represent appealing strategies to create hierarchically-organized hydrogels^[30,31]^. Among “nozzle-free” contactless bioassembly approaches^[31]^, sound-guided assembly offers numerous advantages to transform PAs and fibrin precursors into organized functional composite materials. Sound-guided assembly excels in cytocompatibility and scalability, enabling the simultaneous spatial organization of hundreds to thousands of cells, hydrogel building blocks and inorganic particles at a micrometer-scale resolution precision in a ‘2.5D’ *membrane-like* configurations. By applying mechanical vibrations from a sound source beneath a container of precursor solution, Faraday waves are generated at the liquid–air interface. These waves produce hydrodynamic drag forces that rapidly (within seconds) drive the assembly toward defined positions at the bottom of the liquid^[32]^. Such an approach has been increasingly employed to engineer complex 3D *in vitro* systems, including cardiac tissue constructs^[33]^, spatially-organized capillary networks^[34]^, reproducible nerve ingrowth models for low back pain^[35]^, tumor microenvironment models^[36]^, and architectures that mimic hepatic lobule organization^[37]^. Interestingly, recent advances have extended the use of sound-guided assembly beyond cell aggregation, showing promising potential in the biofabrication of millimeter- to centimeter-scale soft robotic systems^[38]^. Leveraging on the organization of particles within a hydrogel, new opportunities for manufacturing of complex composite/hybrid material can emerge. Indeed, alongside supramolecular assembly operating at the nanoscale, sound-based assembly can act as a micro- and macroscale ‘organizer’ of material building blocks to create fibrin-based composite materials with controlled structural and mechanical properties.

Based on this rationale, in this work we converged supramolecular and sound-based assembly strategies into a ‘one-pot’ workflow to create peptide-protein hydrogels with predictable and controlled mechanical and biological properties. As proof-of-concept, we fabricated and characterized a hybrid hydrogel composite membrane based on the co-assembly of PAs with fibrin into a soft matrix containing sound-patterned stiff calcined bone particles (CBPs) (**Figure 1a**). Our fabrication process spanned multiple scales of order to achieve functionality. At the nano-to-mesoscale, PAs were co-assembled with fibrinogen molecules to create a nanofibrillar mesh able to entrap CBPs into precise patterns. To enrich the fibrin network with precise bioactivity, PAs were designed to show bone morphogenetic protein-2 (BMP-2) binding epitopes (**Figure 1b**), enabling precise BMP-2 binding and release over time. At the meso-to-macroscale, stiff and inorganic CBPs were distributed into organized patterns within PA-fibrin gels, enabling the creation of anisotropic CBP-reinforced PA-fibrin composite membranes. For the first time, a theoretical and experimental relationship was demonstrated between the frequency of applied sound waves and the final mechanical properties of the patterned constructs, providing an innovative design-to-function tool for composite hydrogel fabrication. PA-fibrin hydrogels resulted into more stable materials than normal fibrin and were structurally permissive for cell infiltration. Moreover, by exposing our material to peripheral blood mononuclear cells (PBMCs), we demonstrated how immunomodulatory properties of PA-fibrin hydrogels could be modulated by selection of the additive PAs, unlocking new tools for the design of fibrin-based materials with controlled immunomodulatory properties. This study provides a benchmark for biomaterial design, in which supramolecular and acoustic bioassembly are synergistically exploited to create composite materials with rationally-designed shape-to-function features.

**Figure 1.**
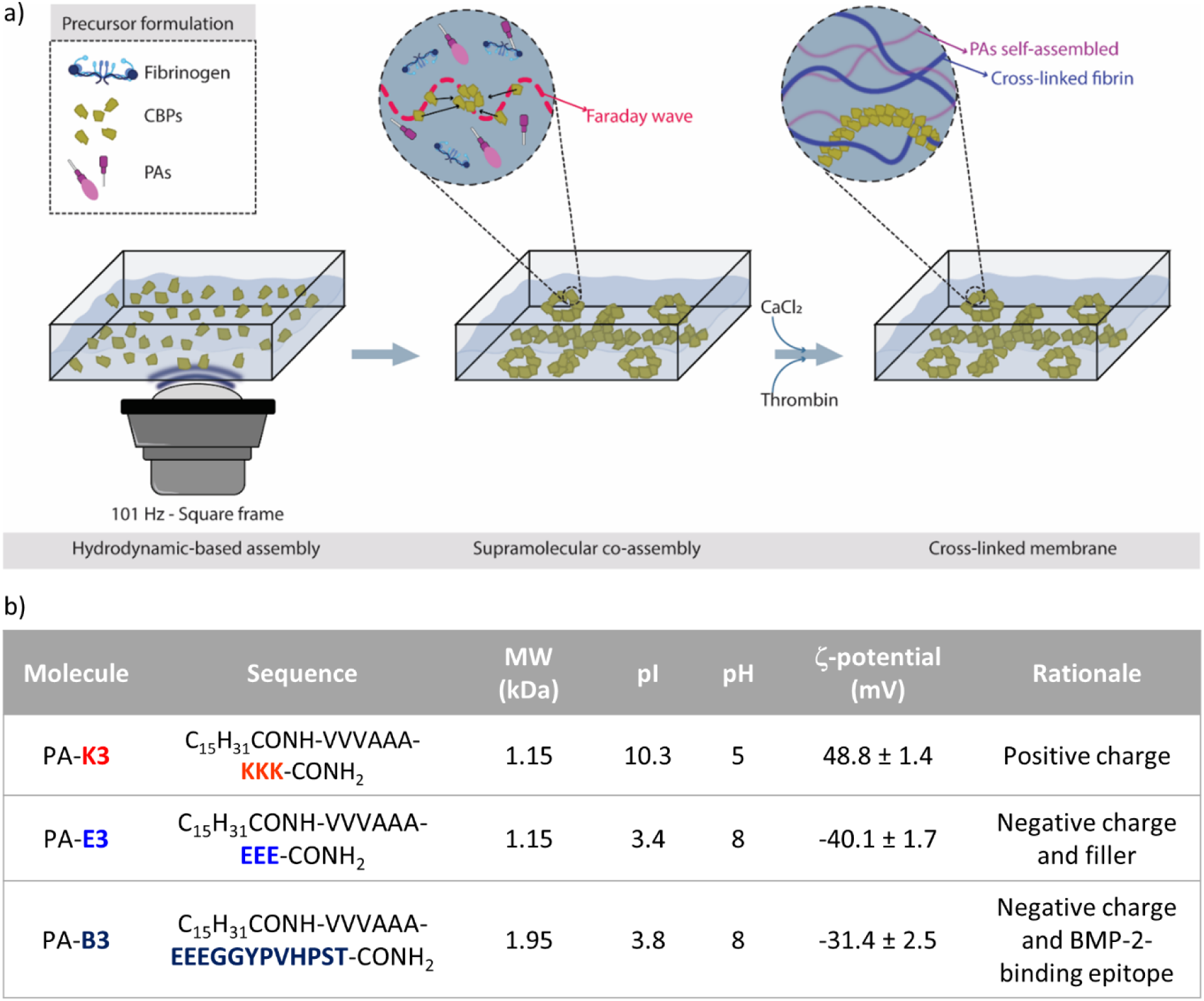
Fabrication strategy of functional hydrogel membranes. a) The schematic illustrates the multiscale approach converging supramolecular co-assembly and sound-guided hydrodynamic assembly to fabricate fibrin-based hybrid hydrogel membranes. From left to right: A precursor solution containing PAs, fibrinogen, and CBPs building blocks is exposed to sound-generated Faraday waves that allows a highly controlled spatial arrangement of CBPs within the precursor volume. The hydrogel solution is then exposed to CaCl_2_ and thrombin that synergistically co-assemble PAs and fibrin into a fibrillar mesh. b) Table summarizing the key physico-chemical information and rationale of the PA molecules used in the study. Data are shown as mean ± standard deviation.

## Results and Discussion

### PA-fibrin co-assembly and characterization

A series of *in vitro* preliminary tests were conducted to assess whether fibrinogen was able to co-assemble with PAs to form self-supporting PA-fibrin hydrogels. To understand the effect of peptide charge on the co-assembly, one negatively charged (PA-**E3**) and one positively charged PA sequence (PA-**K3**) were selected for the study (**Figure 2a**). With the mindset of creating a clinically-relevant peptide-fibrin composite, a clinically approved version of fibrinogen and thrombin, namely EVICEL®, was selected for the experiments. As shown by zeta potential (**Figure 2b**), PA-**K3** was positively charged at neutral pH (ζ_PA-K3_=48.77±1.48 mV), while PA-**E3** carried a negative charge (ζ_PA-E3_=-40.1±1.66 mV). At the same pH, fibrinogen was negatively charged (ζ_Fg_=-6.82±0.24 mV), while thrombin showed an overall net positive charge (ζ_Thr_=7.46±0.45 mV). Due to the charge differences between PA-**E3** and PA-**K3**, it is expected that PA-**E3** and fibrinogen molecules will experience a repulsive force in solution, while an electrostatic complexation is expected to occur between PA-**K3** and fibrinogen. As shown in **Figure 2b**, a mixture of PA-**K3** with fibrinogen led to a drastic reduction of the peptide’s zeta potential (ζ_Fg+PA-K3_=14.53±1.16 mV), while an overall negative charge was preserved when fibrinogen was mixed with PA-**E3** (ζ_Fg+PA-E3_=-7.13±0.53 mV). In both cases, mixtures of PAs and fibrinogen did not produce hydrogels upon contact. Considering the results, it is expected that the final properties of PA-fibrin gels will depend on the charge of selected PAs, as complexation with fibrinogen by PAs could affect the interaction of thrombin with accessible fibrinogen.

**Figure 2.**
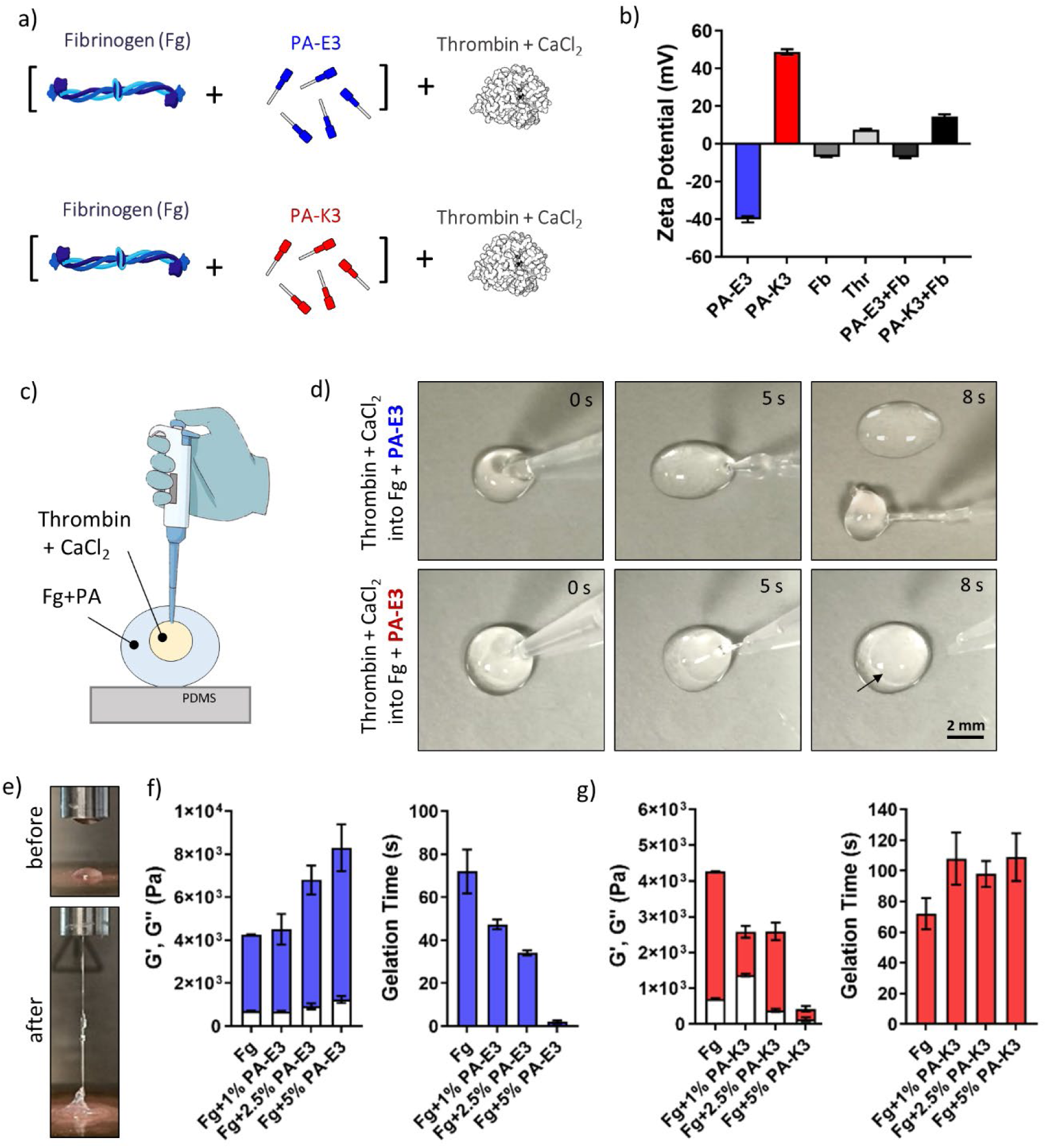
PA-fibrin co-assembly and characterization. a) Schematics of mixture: fibrinogen is pre-mixed with PAs and then mixed with thrombin solution. b) Zeta potential of PAs, fibrinogen, thrombin and mixtures. c) Schematic diagram of PA-fibrin co-assembly. d) Visual assessment of gel formation after injection of thrombin and CaCl_2_ into a droplet of PAs pre-mixed with fibrinogen. e) Rheology set-up and visualization of gel formation. f-i) Gel stiffness and gelation time of fibrinogen mixed with (f-g) PA-E3 and (h-i) PA-K3 after addition of thrombin and CaCl_2._

To test the effect of PAs on the formation of PA-fibrin hybrid gels, a droplet of activated thrombin solution containing CaCl_2_ was injected into a droplet of fibrinogen pre-mixed with PA-**K3** or PA-**E3**, as shown in **Figure 2c**. In this setup, thrombin will initiate the polymerization of fibrinogen into fibrin, while CaCl_2_ will trigger the growth of PA cylindrical micelles into elongated nanofibres. As shown in **Figure 2d**, injection of thrombin and CaCl_2_ in fibrinogen plus PA-**E3** produced an immediate gelation of the solution (<5 s), with the formation of a self-supporting hydrogel able to be collected upon retrieval of the pipette tip. Conversely, a weak gel was generated after mixing thrombin and CaCl_2_ with fibrinogen and PA-**K3**. Moved by this result, a series of oscillatory rheology tests were performed to induce *in situ* PA-fibrin gelation (**Figure 2e**) and assess the stiffness and the rate of gelation of the formed hydrogels. As shown in **Figure 2f**, increasing concentrations of PA-**E3** caused an increase in the stiffness of PA-fibrin gels, whose gelation time was drastically reduced from 72±10.2 s to 2.05±0.74 s upon addition of PA-**E3** (**Figure 2g**). On the other hand, as also observed by visual inspection in **Figure 2d**, addition of PA-**K3** to fibrinogen solutions has a lesser effect on the gelation of PA-**K3**-fibrin. PA-fibrin gels appeared weaker as the content of PAs was increased (**Figure 2h**), while overall PA-**K3**-fibrin gels needed more time to undergo gelation compared to native fibrin (109±15.5 s vs 72±10.2 s), independently of the concentration of PA-**K3** supplied (**Figure 2i**). Moreover, PA-fibrin gels obtained from PA-**E3** were highly stable in PBS over time, with gels being less prone to enzymatic degradation (remaining mass=95.40±1.39% after 7 days) by human metalloproteinase-1, known to have fibrinolytic activity^[39]^, compared to fibrin (mass=81.24±1.94%, **Figure S1**). Coupled together, these results showed that the addition of PA-**E3** to fibrinogen led to PA-fibrin gels with tunable mechanical properties and faster gelation time, as well as higher resistance to enzymatic degradation compared to native fibrin.

### Mechanical and biological functionalization of PA-fibrin gels

Having ascertained that fibrin can be co-assembled with PAs to create tunable PA-fibrin hydrogels, the next step aimed to enrich PA-fibrin with biological and mechanical cues to overcome fibrin’s inherent limitations. Fibrin is known to contain inherent bioactive epitopes, including heparin-binding domains, which enable it to directly sequester and retain growth factors over time; however, the incorporation of additional macromolecules, such as heparin, is essential to effectively sequester and retain growth factors over time^[6]^. In our hybrid fibrin gels design, PAs can play a pivotal role for GF sequestration and release over time, as by designing the chemistry of their hydrophilic head, PAs can self-assemble into nanofibres presenting bioactive function-encoding epitopes on their surfaces. PAs lacking bioactive epitopes, also known as fillers, can be mixed with PAs showing these motifs to create nanofibres with wide ranges of bioactive epitope densities^[40,41]^. In our study, PA-**E3** sequences acted as filler to be mixed with epitopes binding BMP-2 (PA-**B3**, see sequence in **Figure 1b**) to create BMP-2-binding nanofibres (PA-**E3B**). A BMP-2-binding epitope was selected as a proof-of-concept biological functionalization of fibrin, due to the key role played by BMP-2 sequestration and solid-phase presentation for osteogenesis^[42]^. In our design, the heptapeptide sequence ‘YPVHPST’ was used as a BMP-2-binding epitope, due to its high affinity reported in phage display with BMP growth factors (GFs)^[43]^. After assembly, PA-**E3B** nanofibres showed great ability to bind free BMP-2 molecules, as suggested by a significant increase in their molecular size upon contact with growth factor (**Figure 3a**). This BMP-sequestering ability was also evident in 3D constructs, when PA-**E3B**-containing hydrogels were able to physically entrap BMP molecules in their network when exposed to BMP-2-containing solutions (**Figure 3b**). By tuning the epitope density (*i.e.* ratio of PA-**E3** and PA-**E3** displaying YPVHPST) it was possible to tune the amount of BMP-2 released over time (**Figure 3c**), creating a tunable depot for BMP release. Moreover, by selecting different epitope-displaying PAs, it was possible to bind and release alternative GFs from the BMP family, as TGF-β1 using the heptapeptide ‘LPLGNSH’^[20,44]^, making the platform highly modular and versatile (**Figure S2**). Considering these results, PAs can be used to modify fibrin and enhance its limitation to bind and release relevant GFs in a controlled fashion.

**Figure 3.**
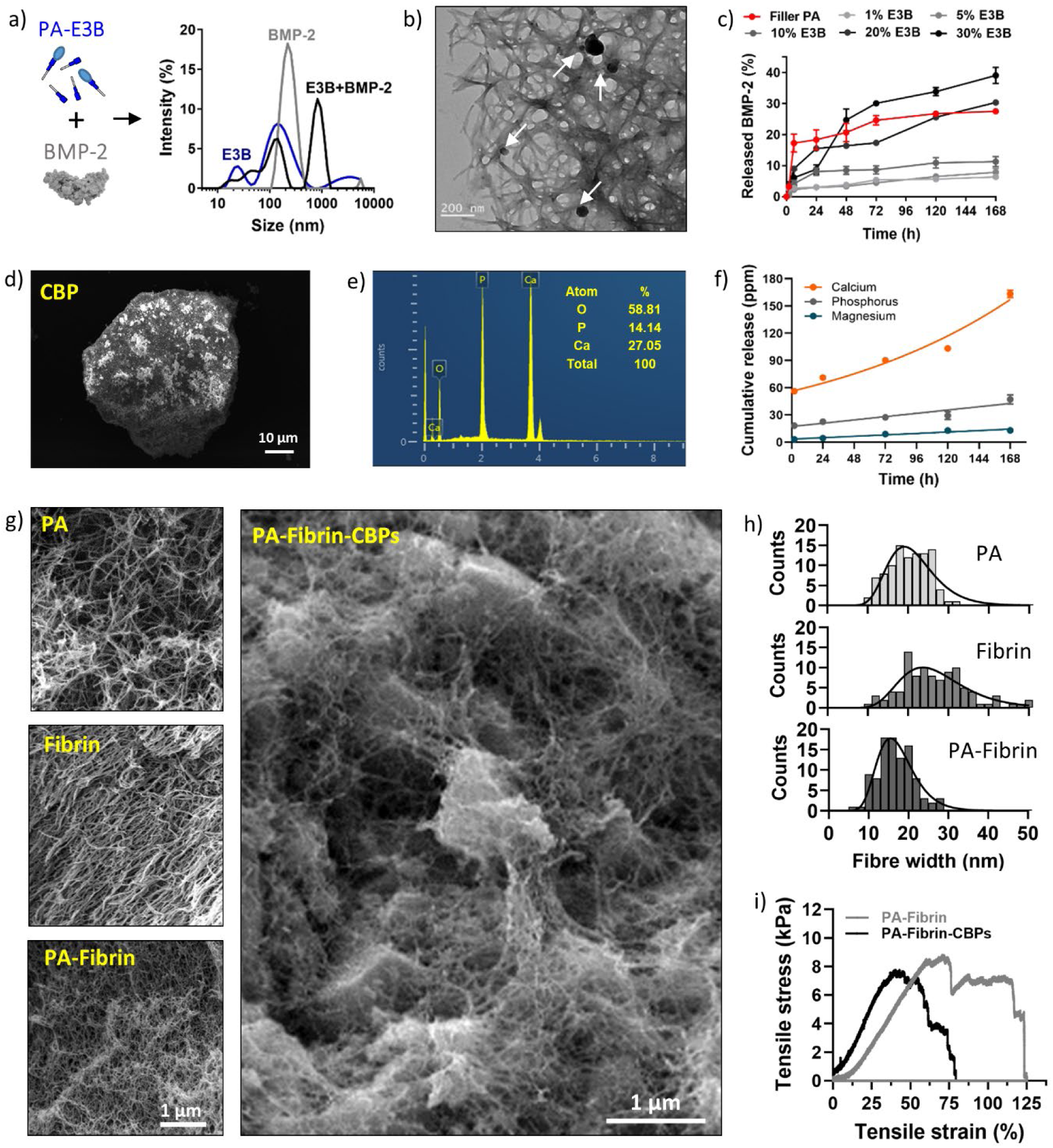
Biological and mechanical functionalization of PA-fibrin hydrogels. a) Size analysis of PA-E3B nanofibres, BMP molecules and PA-E3B after exposure to BMP-2. b) BMP-2 molecules (black features indicated by white arrows) entrapped in PA-E3B hydrogels. c) ELISA assays of released BMP-2 from PA-E3B gels show dependence of release profiles on epitope density. d) SEM image of a CBP. e) EDX analysis of a CBP. f) Cumulative release of calcium, phosphorus and magnesium ions from CBPs dispersed in PBS and measured by ICP-OES. g) SEM images of PA, fibrin, PA-fibrin and PA-fibrin-CBP hydrogel networks with h) their relative fibre diameter distribution. i) Uniaxial load tensile test of PA-fibrin hydrogel membranes with and without addition of CBPs.

Alongside bioactivity, fibrin is also known to have poor mechanical properties. Moreover, for bone applications, fibrin alone is known to have poor osteoconductive properties, not able to initiate nor support biomineralization. To overcome these limitations, we explored the possibility to incorporate CBPs within PA-fibrin constructs. CBPs are polygonal microparticles obtained from calcination and milling of bones to obtain powders highly rich in inorganic mineral components, mainly hydroxyapatite^[45,46]^ (**Figure 3d**). Exploiting our experience with the fabrication of CBPs, we produced in house CBPs with a distribution size of 80-180 µm (**Figure S3**), within the range of particle size previously used by our group in other sound-driven bio-assemblies^[32,34]^. For our CBPs, we confirmed the presence of a high content of calcium phosphates products, with a Ca/P ratio of ∼1.91, ascribable to mineral structures containing calcium-rich (*e.g.* CaO) or tricalcium phosphates (TCP) phases^[47]^ (**Figure 3e**). These phases resulted highly dynamic when dispersed in solution, with release of calcium and phosphorus ions over time, as observed *via* ICP-OES analysis (**Figure 3f**). To create reinforced PA-fibrin hydrogel membranes, CBPs were physically dispersed into PA-fibrinogen solutions and then entrapped into PA-fibrin gels upon addition of activated thrombin solution containing CaCl_2_. The process of PA-fibrin co-assembly did not create a visible phase separation between PA-based and fibrin-based networks, instead the two building blocks synergistically created a hybrid nanofibrillar network with unimodal fibre diameter distribution, as occurs for pure PA and fibrin gels before co-assembly (**Figure 3g**). This feature is quite appealing for the generation of fibrin-based materials, as phase separation often represents one of biggest drawbacks for hybrid fibrin-based hybrid materials^[4]^. Interestingly, the inclusion of PAs induced the formation of a nanofibrillar hybrid fibrin with thinner fibres (µ_PA-Fib_=16.76±1.32 nm) compared to normal fibrin (µ_Fib_=26.1±1.37 nm), possibly due to PAs positioning affecting lateral aggregation and width growth of fibrin protofibrils (**Figure 3h**). The same fibre diameters were preserved upon addition of CBPs (µ_PA-Fib+CBPs_=17.53±1.23 nm). After gelation of the whole composite, CBPs resulted evenly distributed and mechanically interlocked within the PA-fibrin nanofibrillar mesh, as shown in **Figure 2g**. In terms of mechanics, the addition of CBPs resulted in a significant increase of stiffness for PA-fibrin-CBPs (*E=*21.53±1.90 kPa) compared with CBP-free counterparts (*E=*18.83±3.25 kPa, **Figure 2i**). Coupled together, these results showed that fibrin can be ‘enhanced’ both biologically and mechanically by introduction of bioactive PA sequences and dispersion of inorganic microparticles, creating fibrin-based materials with higher stiffness and enhanced growth factor-binding abilities.

### Fabrication of functional hydrogel membranes via sound-guided assembly

After hydrogel membrane functionalization via inclusion of bioactive epitopes and physical entrapment of CBPs, our next step of material engineering exploited sound-guided assembly to create functional hydrogel membranes with dictated microstructures. In our previous works, we harnessed sound-guided Faraday waves and their drag forces to precisely spatially arrange microparticles or cellular aggregates within hydrogel precursors, including gelatin methacryloyl (GelMA)^[32]^, collagen^[35]^ and fibrin^[36]^. This fabrication method features a cost-effective, bench-top wave generator, on top of which a petri dish containing the liquid precursor formulation is securely positioned (**Figure 1a**). When activated and tuned with a specific frequency and amplitude, the wave generator allows the formation of Faraday waves onto the precursor surface, thus inducing drag forces within the bulk volume. This physical phenomenon allows objects such as nano- and micro-particles, single cells, or cell aggregates, to be spatially arranged within the hydrogel precursor into specific configurations that are highly dependent on the acoustic frequency and the container shape. Based on this rationale, a wide range of frequencies (0 – 143 Hz) was tested using a container featuring either a circular or a square geometry. As shown in **Figure 4a**, by tuning these few parameters, distinct patterns were generated within the liquid hydrogel precursor, eventually creating different soft nanocomposites in which the fillers (*e.g.*, CBPs) featured a distinct spatial arrangement within their matrices (*e.g.*, PA-fibrinogen solution). At lower frequencies (0-57 Hz), larger and looser patterns were obtained, while higher frequencies (75-143 Hz) led to finer and closer-packed patterns. Numerical simulations of the surface displacements of the hydrogel precursor (**Figure 4b, Video S1**) and simulations of the hydrodynamic drag forces generated within highlighted a tendency of the CBPs to cluster in correspondence with areas featuring the lowest force potential, thus characterized by limited turbulence (**Figure 4c**). At 101Hz, the areas characterized by the lowest drag forces, corresponded to a pattern featuring four points of symmetry and a central equal-armed cross surrounded by four rings (**Figure 4d**). The comparison between the experimental data (**Figure 4a**) and the *in silico* simulations (**Figure 4b-d**) further strengthened the role of container shape and acoustic frequency as key orchestrators of the sound-driven pattern formation, with a bidirectional relationship between the computer simulations and the laboratory experiments. The predictability and tunability of the patterns was consistent with previous reports on acoustic-driven assembly, from our lab, although processing other hydrogel precursors. At this stage of material design, *in silico* simulations were essential to understand how the acoustic waves generated by the sound-guided assembly would interact with the hydrogel precursor, and with its embedded CBPs to guide the formation of specific CBP patterns within PA-fibrin matrices. The relationship between acoustic frequency and CBP patterns within PA-fibrin matrices was further assessed by analyzing the trend of the pattern, coastlines and areas, as a function of the frequency.

**Figure 4.**
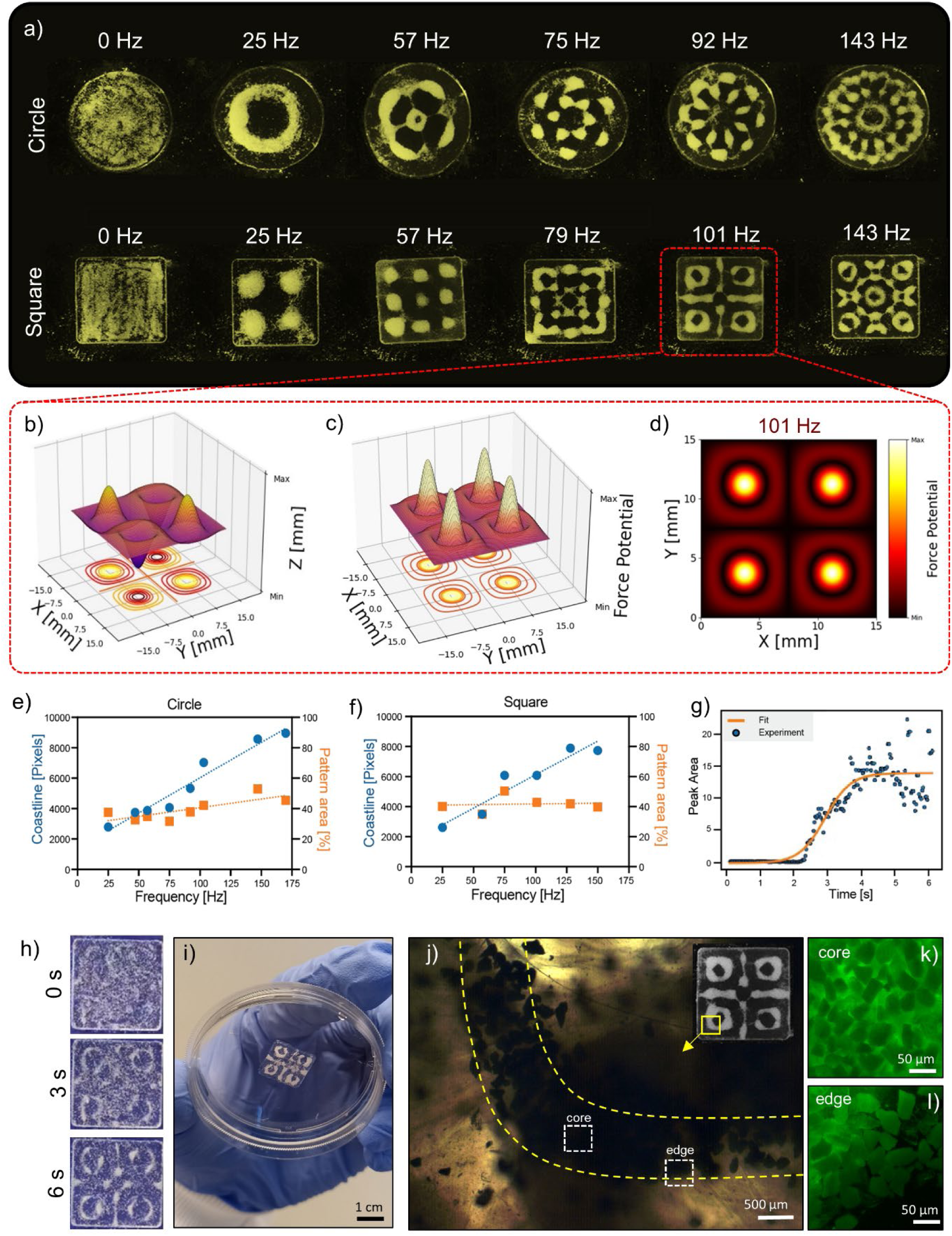
Sound-guided assembly of functional PA-fibrin-CBP membranes. a) CBP patterns obtained within PA-fibrinogen solutions contained in circle and square containers. b) Surface displacement, c) drag force potential plot and d) its cross-section for a PA-fibrinogen solution excited with sound waves of 101Hz. e) Relationships between pattern coastlines and areas with applied acoustic frequency for e) circle and f) square containers. g) Kinetics of pattern formation. h) Time frames of the CBP pattern formed at 101Hz over time. i) PA-fibrin-CBP hydrogel membrane with pattern induced at 101 Hz. j) Detail of the pattern at 101Hz (quarter of the top left corner ring) under polarised light microscopy. Details on CBP packing at k) core and l) edge of the pattern.

Figure 4e-f showed a linear correlation between the increment of the pattern coastlines and the acoustic frequency, while this trend was minimal for the pattern area, regardless of the container shape. We could speculate that an increase in acoustic frequency would lead to smaller wavelengths (nearly punctual for infinite frequencies) that are translated into more densely packed CBP patterns. Conversely, low pitch sound waves with larger wavelengths induce looser patterns of CBPs, as observed for frequencies below 60 Hz. This patterning behavior followed the physics of Faraday wave generation, where higher frequencies lead to shorter wavelengths and more compact structures. Overall, the process of sound-guided bio-assembly occurred within tens of seconds, with patterns becoming stable approximately four seconds after the application of acoustic waves (Figure 4g). Here, experimental data of pattern kinetics followed theoretical expectations, further confirming the reproducibility and predictability of the bio-assembly process (Figure 4g), in agreement with previous kinetics results observed with GelMA matrices^[32]^. Once CBP patterns were formed within PA-fibrinogen solutions within seconds (Figure 4h**, Video S2**), these were entrapped in PA-fibrin gels by the addition of thrombin and CaCl_2,_ which elongate PA nanofibres and enables fibrin polymerization. Sound-patterned PA-fibrin-CBP were self-supporting upon gelation, as shown in Figure 4i. It is worth mentioning that PA-fibrinogen solutions represent an ideal precursor for sound-guided assembly for two main reasons. The first one relies on the low viscosity, time-dependent and shear-thinning behavior of PA-fibrinogen solutions, able to modulate and reduce its viscosity upon application of high shear rates, even in cyclic states of shear simulating sinusoidal sound waves (**Figure S4**). Second, contrary to how collagen or gelatin behave, for example, here gelation was induced enzymatically and chemically (*via* thrombin and CaCl_2_), with no need for temperature switches as gelation triggers. This feature is highly appealing for material fabrication, as it removes the challenge of dealing with thermo-sensitive precursors, which are often accompanied with shape distortions, such as contraction, just after patterning.

Looking at the microscale, we could observe how sound waves and Faraday wave drag forces were able to ‘sculpt’ the PA-fibrin hydrogel networks. Indeed, under polarized light, distinct anisotropic patterns within the PA-fibrin matrix were observed, with birefringent lines following the curvature of the induced pattern (Figure 4j). Similar alignments of peptide nanofibres under the effect of sound waves have been observed by Ulijn and colleagues, who showed the formation of anisotropic structures at the micrometer scale within tripeptide hydrogels subjected to oscillating pressure waves^[48]^. On a similar note, Hocking and Dalecki observed micrometric deformation of collagen gels fabricated in the presence of ultrasound^[49,50]^. By pre-staining CBPs with calcein, we were also able to appreciate the effect of sound waves on the organization of CBPs at the microscale. By looking at the core area of the pattern, we observed densely-packed CBPs configurations, adopting a maximum packing density in the core of the patterned structures (Figure 4k). Conversely, by moving towards the edges of the pattern, it was visible how CBPs were more loosely organized, with a certain level of inter-particle distance that seemed to increase moving away from the core of the patterns (Figure 4l). This observation provides a visual representation of the effect of the Faraday drag forces on CBPs, which become densely packed at the local minima of the drag force potential (core of the patterns) and loosely arranged moving away from those minima (edges of the patterns). Altogether, these observations highlighted how acoustic frequency can be used to ‘sculpt’ hydrogel membranes with high level at the microscale.

### Sound-induced patterns dictate the mechanics of hydrogel membranes

Bioconvergence of supramolecular and sound-guided assembly enabled the fabrication of composite hydrogel membranes with high compliance and elasticity. Samples could be stretched multiple times without any delamination of the embedded CBPs from the peptide-fibrin mesh, alongside without any visible loss of the sound-induced pattern (Figure 5a**, Video S3**). As the configuration of fillers within a nanocomposite is highly correlated with its bulk mechanical properties, we speculated that for our composite hydrogels, the configuration of CBPs within their PA-**E3B**-fibrin matrix could also affect the final mechanical properties of the composite material. In particular, due to the possibility of using Faraday waves as organizers of CBPs and ‘sculptors’ of hierarchical structures, we hypothesized that finer patterns obtained at higher sound wave frequencies could exhibit higher mechanical strength compared to coarser ones obtained at lower frequencies. To prove this hypothesis, we ran uniaxial tensile tests on fabricated hydrogel membranes and the stress vs strain curves are presented in Figure 5b. All membranes showed a linear elastic region followed by a plastic behavior, as expected for viscoelastic materials. Analyses of the Young’s moduli showed that at 0 Hz, when no sound was applied and when CBPs were randomly organized within the PA-fibrin matrix, hydrogel membranes showed a modulus of *E_0Hz_*=21.53±1.90 kPa, while upon application of 25 Hz and formation of the first pattern, the Young’s modulus experienced a drastic drop to *E_25Hz_=*5.80±1.01 kPa (Figure 5c). This ‘softening’ effect was expected, as stiffness for composite hydrogels highly depend on the organization of fillers within the matrix and how this propagates cracks upon application of a tensile stress. Indeed, increasing frequencies of sound waves induced finer patterns showing shorter distances between CBPs, allowing higher resistance to crack propagation and thus stiffer hydrogel membranes. For instance, patterns created at 101 Hz induced a membrane stiffness of *E_101Hz_=*14.62±1.25 kPa, while at 143Hz we reported a stiffness of *E_101Hz_=*17.69±2.36 kPa. Overall, from the tensile tests, a linear relationship emerged between the frequency applied to form the patterns of CBPs within the PA-fibrin matrix and the bulk stiffness of the PA-fibrin-CBPs composite material.

**Figure 5.**
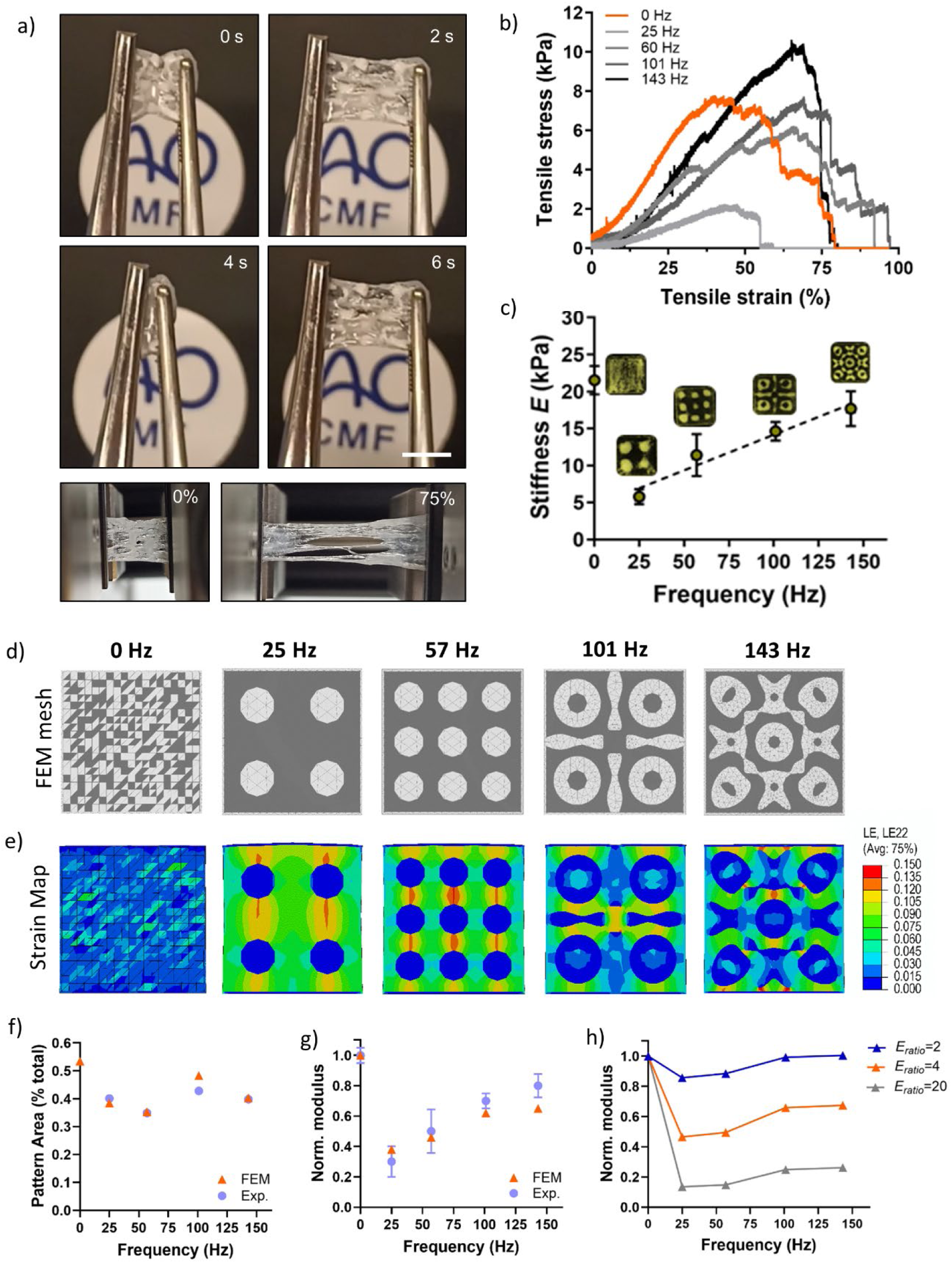
Experimental and in silico mechanical tests. a) Time-frames of PA-fibrin-CBP membranes stretched multiple times with tweezers (above) and the same membranes at 0% and 75% of strain deformations. b) Tensile tests of PA-fibrin-CBP membranes and c) Young’s moduli vs frequency used to induce the sound-based patterns. d) FEM meshes for the five models generated where the subdivision of the two phases is visible (gray = PA-fibrin hydrogel, white = CBPs), and e) logarithmic strain maps in the loading direction at the same maximum loading (linear regime) from FEM simulations of patterned membranes under uniaxial tension. f) Comparison of experimental and FEM measurements of pattern area, expressed as a proportion of the total area of the square membrane. g) Comparison between experimental and simulated Young’s moduli (normalized by the 0 Hz modulus). h) Sensitivity analysis showing effective Young’s modulus as a function of E_ratio_ for representative patterned geometries.

Intrigued by this behavior, we questioned whether these observations could be generalized and modelled *in silico* to create a toolkit where the stiffness of acoustically-patterned composite hydrogels could be predicted starting from the frequency of the sound waves. A similar approach has been applied for the bioplotting of nonapeptide hydrogels, where printing results were modelled starting from the hydrogel viscosities^[51]^. In this case, finite element method (FEM) simulations were performed to investigate the mechanical response of acoustically-patterned PA-fibrin membranes containing a CBP phase. The primary objective was then to validate whether the patterns generated by acoustic assembly could be used to control and predict the mechanical properties of the resulting composite membranes. In particular, this modeling effort supports the hypothesis that the patterning process enables tunable mechanics based on controlled spatial organization of different phases. Starting from the experimental images of the fabricated composite hydrogels, the geometries for the FEM simulations were derived for the frequencies of interest and meshed (Figure 5d) using linear triangular finite elements for mechanical simulations. Figure 5e shows the internal strain distributions of the composite hydrogel under tension and how the local deformations across the areas of the hydrogels highly depend on the pattern induced. Stress maps during the loading process up to the maximum value simulated are shown in **Video S4-S6**. Low frequencies (*e.g.* 25 and 57 Hz) induced high deformations in the matrix regions bridging CBP agglomerates, while higher frequencies (*e.g.* 101 and 143 Hz) showed a more even distribution of local deformation with lower magnitudes. As observed previously in Figure 4f, also the FEM analysis showed that experimental pattern area was constant across frequencies and could not explain the stiffness variation (Figure 5f). Conversely, a linear relationship between the frequency applied to obtain the CBP pattern and the stiffness of the PA-fibrin-CBP hydrogel was observed, which was in line with what observed experimentally (Figure 5g). Interestingly, by tuning the ratio of Young’s moduli of PA-fibrin and CBP phases (*E_ratio_*), it was possible to correlate its influence on the global stiffness of the composite (Figure 5h). Overall, as computed by the FEM analysis, the stiffness of composite hydrogels was still dependent on the frequency of waves applied, with *E_ratio_* inversely proportional to the bulk stiffness. This analysis suggested that the mechanical tunability experimentally observed is consistent with physically-plausible variations in the stiffness of the membrane filler. These simulations demonstrated that the mechanical properties of acoustically-patterned PA-fibrin-CBP membranes can be effectively predicted *via* FEM, with geometry and material contrasts extracted directly from experimental data. We envisage that this ‘frequency-stiffness’ relationship could be generalized and applied for the fabrication of alternative sound-assembled composites, still based on the acoustic patterning of ‘stiff’ particles (*e.g.* organoids, ceramics, glass) within softer matrices of precursors (*e.g.* Matrigel, resins), for wide applicability in the healthcare (*e.g.* organ-on-a-chips, bone plates) and advanced manufacturing sectors (*e.g.* composite aircrafts).

Supramolecular and sound-induced assembly can dictate cell organisation and response Once the bulk properties were characterized and the fabrication steps were established, the following step focused on assessing whether supramolecular and sound-induced assemblies could be exploited to dictate cell organisation and response. Towards this goal, first mesenchymal stromal cell (MSCs) spheroids were dispersed in the PA-**E3B**-fibrinogen solution and sound was applied as accomplished previously for CBPs (Figure 6a). Interestingly, MSC spheroids with a nominal diameter of ∼50 μm and larger CBPs (Figure 6b**, Video S7**, and Figure 4h**, Video S2**, 101 Hz) were acoustically organized, via Faraday waves, into the same pattern shape. While the spatial resolution of such hydrodynamic assemblies is generally influenced by particle size, in this case both MSC spheroids and CBPs fall within a size range that enables their organisation into the same final structure at a frequency of 101 Hz. This observation further confirms that the final pattern can be predicted *a priori* through numerical simulations, as demonstrated in Figure 4. These results also extended our library of ‘sound-patternable’ materials, adding PA-fibrinogen alongside the already explored GelMA^[32]^, collagen^[35]^ and pure fibrin^[36]^. Building in complexity, in our previous work we also explored the use of both MSC spheroids and β-TCP microparticles as objects to be sound-patterned for the creation of anisotropic hydrogels. When both MSCs and TCPs were exposed to sound waves, a dual distribution of cells and particles emerged, due to the differences in the size and density of the two objects^[52]^. A dual distribution has been observed by Pu Chen and colleagues, who assembled induced pluripotent stem cell-derived liver spheroids and endothelial cells into hexagonal cytoarchitectures using acoustic differential bioassembly^[37]^. Starting from this, we questioned what could happen to MSC seeded on hydrogels containing CBPs already patterned. As shown in Figure 6b, MSCs started to penetrate into the PA-fibrin-CBP hydrogel membranes, reaching the bottom of the gels within 48 hours (Figure 6c). During this timeframe, penetrating MSCs adopted a prominent dual distribution with the CBPs over time, with cells organising either within or surrounding the patterned particles (Figure 6d-g). This spatial organisation of MSCs/CBPs resembled the dual distribution obtained for MSCs and microparticles embedded in the liquid precursors and subjected to sound waves^[52]^, suggesting identical patterning either with cells used as objects to be patterned or with cells deposited on the already patterned hydrogels. This observation points to the fact that cell distribution is dependent on the microarchitectures that sound waves have formed, possibly due to hydrogel network densifying in specific points forcing cells to organise into specific configurations. This was further supported by radial colocalization analysis, performed on four independent pattern regions (N = 4) within each hydrogel. Despite slight variability among regions, the profiles showed a consistent trend, with MSCs progressively accumulating in the surroundings or in between the areas of the patterned CBPs over time. This convergence in spatial distribution across multiple pattern nodes confirms that the observed dual localization is a reproducible outcome of the underlying patterning, rather than a local anomaly. Thus, acoustic cues serve as initial cell guidance templates, dictating cell organisation and migration. MSCs penetrating within the patterned hydrogels remained highly viable with sustained metabolic activity, as confirmed by Live/dead and Presto Blue® assays (Figure 6h-i). Collectively, these results suggest that microscale patterning can guide cell penetration and organisation acting as microstructrural cues that cells leverage for spatial organisation.

**Figure 6.**
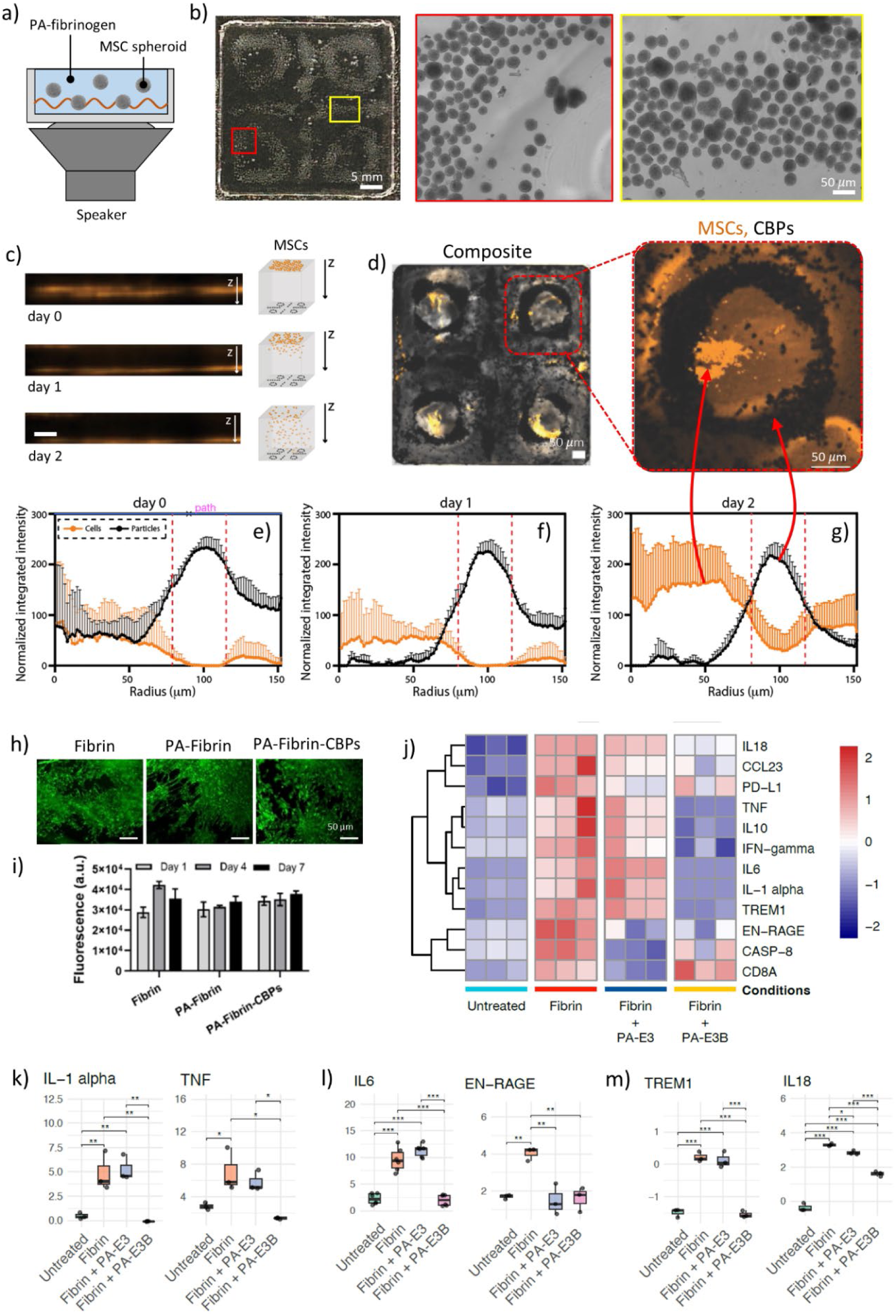
Supramolecular and sound assembly dictate cell organisation and response. (a) Schematics of sound-patterning of MSC spheroids into a PA-fibrinogen bath. b) Formation of a spheroid pattern and details of cell aggregation. (c) Confocal z-stack reconstructions illustrating vertical migration of hMSCs through PA-fibrin hydrogels over time. (d) Overlay of PKH-labelled hMSC membranes (orange) and CBPs (dark spots) confirming co-localization at nodal regions. Data are expressed as mean ± SD across four distinct patterned regions (n = 4 ROIs per time point) within the same hydrogel membrane. (e–g) Radial distribution profiles of hMSCs (black) and CBPs (orange) at day 0, 1, and 2, respectively, showing divergence in migration patterns. (h) Live-cell staining of hMSCs seeded on Fibrin, PA-**E3B**-fibrin, and PA-**E3B**-fibrin-CBPs scaffolds. (i) Quantification of cell proliferation via PrestoBlue® metabolic activity assays over 7 days (n = 5). (j) Proteomics heat-map using the Olink Proximity Extension Assay to explore immune and inflammatory responses of PBMCs seeded on culture plastic, fibrin, PA-**E3**-fibrin and PA-**E3B**-fibrin hydrogels. Bar plots of selected markers: k) IL-1α, TNF; l) IL-6, EN-RAGE; m) TREM1 and IL-18.

As a last step of investigation, we tested the possibility of tuning the bioactivity of PA-fibrin gels by choosing different epitope-displaying PAs. To do this, composite hydrogels were exposed to peripheral blood mononuclear cells (PBMCs), often used to get an insight into the immunomodulatory surface properties of hydrogels and biomaterials^[53,54]^. In particular, we were interested to see whether the inclusion of different PAs within PA-fibrin hydrogels could be used as a ‘modulator’ of immune cell response, due to the interplay between biomaterials design and activation of the immune system^[55]^. Towards this goal, targeted proteomic analysis was performed after exposure of PBMCs to the fabricated PA-fibrin gels. Overall, the proteomics analysis revealed distinct alterations in inflammation-related proteins across the experimental conditions. As depicted in the heat-map (Figure 6j), fibrin alone induced a marked upregulation of several inflammatory markers, including IL-1α, IL-6, TNF, EN-RAGE, TREM1, and IL-18, compared to the untreated group. Notably, exposure of PBMCs to both PA-**E3**-fibrin and PA-**E3B**-fibrin hydrogels substantially attenuated this fibrin-induced inflammatory response. Among the hybrid fibrin hydrogels tested, PA-**E3B**-fibrin gels demonstrated a slightly stronger anti-inflammatory effect, resulting in significant reductions in key early-response pro-inflammatory markers, such as IL-1α, TNF (Figure 6k), acute-phase response as IL-6, and inflammation modulator markers, such as EN-RAGE, a DAMP molecule often involved as inflammation amplifier via RAGE receptor signaling^[56]^ (Figure 6l). Separate bar plots illustrating IL-18 and TREM1 markers involved as innate immunity coordinators, further confirmed that the introduction of a BMP-2 binding epitope as co-assembly building block within fibrin matrices enabled expression of these markers closer to the baseline levels observed in untreated samples (Figure 6m), with a better anti-inflammatory effect compared to fibrin co-assembled with PA-**E3** and fibrin alone. Collectively, these findings indicate that PA-**E3** and PA-**E3B** mitigate fibrin-induced inflammation in PBMCs, with PA-E3B exhibiting a more pronounced effect. In terms of materials design, these results suggest that by fine choice of fibrin ‘modifiers’, PA-**E3** and PA-**E3B**, it is possible to fabricate hybrid fibrin-based materials with a dictated immunomodulatory profile.

## Conclusions

In summary, we have demonstrated the possibility to biofabricate composite protein-based hydrogel scaffolds with dictated structure-to-function features by exploiting the bio-convergence of two bottom-up fabrication strategies, namely peptide self-assembly and sound-guided bio-assembly. In particular, we have shown how by starting from fibrin hydrogels it was possible to ‘enrich’ these materials both mechanically and biochemically by physical entrapment of inorganic microparticles and by chemical incorporation of bioactive functional groups. In details, by leveraging on the co-assembly of proteins with epitope-displaying PAs, we demonstrated the opportunity to create PA-fibrin hydrogels able to bind and release growth factors, such as BMP-2, for enhanced biological activities, surpassing inherent limitations of traditional fibrin scaffolds. On the other side, by exploiting the use of sound waves to biofabricate composite scaffolds whose fillers are sound-patterned, we embedded inorganic particles, i.e. CBPs within PA-fibrin hydrogels and we were able to pattern their distribution in final hydrodynamically-organized PA-fibrin-CBPs. By merging experimental and *in silico* mechanical tests, we observed that the frequency of the sound waves explored has a direct relationship to the stiffness of the final composite hydrogels, unlocking a new way to create mechanically-robust peptide-protein-based scaffolds. Finally, we demonstrated how these new peptide-protein systems can show biological potential *via in vitro* studies and proteomics studies. Based on our results, we believe we have developed a new biofabrication tool in which supramolecular and sound-based assemblies can be used to create multiple peptide-protein materials with dictated structural, mechanical and translational potential. Here, we presented fibrin-based hydrogel composites, but we envisage that a generalization of this concept is possible for other PAs and protein building block combinations. For instance, more complex mixture of proteins and biofluids, such as blood and bone marrow aspirate^[28]^, could be used as protein-rich phases to co-assemble with GF-sequestering PAs, to create ‘bio-cooperative’ regenerative materials with synergistic properties arising by the means of sound and molecular co-assembly.

## Experimental Section

### Materials

PAs in lyophilised form were purchased from Peptide Synthetics, UK. In particular, two peptides sequences were synthesized, namely PA-E3 with sequence Palmitoyl-VVVAAAEEE and PA-B3 with sequence Palmitoyl-VVVAAAEEE-GGYPVHPST. All received peptides were purified by HPLC and purity values >95% were used for the study. Human fibrin was purchased as EVICEL® kit (Ethicon, UK) containing one vial of human clottable fibrinogen (55-85 mg/mL) and one vial of human thrombin (800-1200 IU/mL). For this study, fibrinogen was diluted two-fold by mixing 1:1 with 1X PBS. Calcined bone particles were produced in-house and kindly supplied by Prof Zhiyu Zhou. If not stated otherwise, all reagents were purchased from Sigma Aldrich, UK.

### PA-fibrin hydrogel formulations

To create PA-fibrin hydrogels, 1% PA solutions were first created and then mixed with fibrinogen solutions before addition of thrombin. Specifically, for PA-**E3B**-fibrin hydrogels, 0.9 mg of lyophilized PA-**E3** powder was mixed with 0.1 mg of lyophilized PA-**B3** powder and vortexed for 20 sec to obtain 1 mg of a powder mixture, namely PA-**E3B**. This mixture was then dissolved in 90 μL of 10 mM HEPES (pH 7.4) and basified with 10 μL of 0.5 M NaOH to create 1% PA-**E3B** solution. Once the PA-**E3B** solution was made, 17 μL of this was injected into 53μL of fibrinogen solution and mixed thoroughly to form PA-**E3B**-fibrinogen solutions. Then, 30 μL of thrombin as added to the mixture to induce gelation and produce 100 μL of PA-**E3B**-fibrin hydrogel. For the PA-**E3**-fibrin and PA-**K3**-fibrin hydrogels, all the steps mentioned above were followed but 1 mg of either PA-**E3** or PA-**K3** lyophilized powder was used.

### ζ-potential and Dynamic Light Scattering (DLS)

ζ-potential was performed to measure the surface charge of PAs, fibrinogen, thrombin and their mixtures. DLS was used to measure the size of BMP-2 molecules (1 µg/mL) and 1% PA-**E3B** after incubation. For both techniques, PAs or proteins of interest were diluted into their solvents at a concentration of 0.1% w/v and solutions were double-filtered with a 0.22 µm syringe-filter before any measurement. ζ-potential and DLS measurements were acquired at 25°C in triplicate on three independent samples using a Zetasizer Nano-ZS ZEN 3600 (Malvern Panalytical, UK).

### Oscillatory rheology

Oscillatory rheology was performed to track the gelation and check the rheological properties of PA-fibrin gels. All rheological measurements were performed using an Anton Paar MCR102 rheometer (St Albans, UK) using a 8 mm parallel plate geometry and a 0.5 mm gap. To check the gelation time of PA-fibrin gels, 70µL of PA-fibrinogen solution was placed on the bottom plate of the rheometer and 30 µL of thrombin was positioned on the surface of the 8mm plate head. The upper head was lowered to allow mixing of thrombin with PA-fibrinogen and a time sweep test was performed immediately. For the time sweep, 0.2% strain and 1Hz frequency values were used. To check the stiffness of PA-fibrin hydrogels, samples were prepared as described earlier and placed onto the rheometer’s static bottom plate while the upper plate was moved to the measuring gap. To measure storage (G’) and loss (G’’) moduli an amplitude sweep test was performed at 37°C and 1 Hz in the 0.01 – 100% strain range.

### Quantification of BMP-2 release *via* ELISA

To assess the release of BMP-2 molecules from hydrogels, 100µL of gels were exposed to 200µL solutions of 1 µg/mL BMP-2 solutions (Peprotech, UK) and at each time point (0, 4, 24, 48, 72, 120 and 168 h) the whole solution in the well was collected and replaced with fresh BMP-2 solution. The amount of BMP-2 in each aliquot was quantified using a BMP-2 DuoSet sandwich ELISA assay (code: DY355, R&D Systems, UK). For this test, six increasing concentrations of BMP-2 binding epitopes were tested by physically mixing PA-**E3** with PA-**B3** powder to get 0% **E3B** (*i.e.* 100% PA-**E3** or ‘Filler PA’), 1%, 5%, 10%, 20% and 30% PA-**E3B**. The cumulative release data points were averaged across three independent replicas and plotted using GraphPad Prism 10 software.

### Transmission Electron Microscopy (TEM)

TEM was used to assess the nanostructure of the hydrogels in the presence of BMP-2 molecules. Hydrogels were diluted 20-fold with ultrapure water and 20µL was pipetted on carbon-coated copper TEM grids (EM Resolutions Ltd, UK) for 30 seconds. Excess solution was carefully removed using Whatman No.1 filter paper and grids were stained with 2 µL of 2% uranyl acetate for 30 s before washing with 18 µL of ultrapure water. After staining, excess water was removed with filter paper and samples were left to air dry for 1 hour. Imaging was performed on a FEI Tecnai Biotwin-12 TEM running at 100 kV and equipped with a Gatan SIS Megaview IV camera.

### Scanning Electron Microscopy (SEM)

SEM was used to investigate the microstructure of PA-**E3B** and fibrin gels, PA-**E3B**-fibrin and PA-**E3B**-fibrin-CBP composite gels. Moreover, SEM was also used to image CBPs and perform EDX elemental analysis. For the hydrogels, samples were prepared as described above and fixed at room temperature with 4% paraformaldehyde solution (PFA) for 20 min. Excess of PFA was removed, sample were washed twice with deionised water and dehydrated with a series of increasing ethanol concentrations (20, 30, 50, 70, 90, 96, and 100%) for 10 min each. Samples were subjected to critical point drying (Leica EM CPD300, Leica, Wetzlar, Germany), mounted on carbon-coated SEM stubs and sputter-coated with iridium (10 nm thick coating) before imaging with an JEOL 7000F FEG-SEM (JEOL Ltd., Tokyo, Japan). Fiber diamter analysis was performed by measuring randomly the lengths of 100 fibres per sample across three independent SEM images using ImageJ software (v1.54j). For the CBP, particles were directly mounted on carbon-coated SEM stubs and sputter-coated with iridium (10 nm thick). EDX analysis was performed with a JEOL IT-200 (JEOL Ltd., Tokyo, Japan).

### Inductively coupled plasma optical emission spectroscopy (ICP-OES)

To test the stability of CBPs, particles 10 mg of particles were first dispersed into 1 mL of 1X PBS and incubated over time at 37°C. At each time point (0, 24, 72, 120, 168 h), CBPs were centrifuged at 4000 rpm for 5mins and supernatant was collected for ICP-OES analysis to reveal the amount of ions released. For this test, calcium (Ca), phosphorus (P) and magnesium (Mg) were investigated as ions of interest. ICP-OES was run on a Perkin Elmer Optima 2000 DV ICP-OES with a cyclonic spray chamber, 2.0 mm alumina injector, double monochromator, and a dual backside-illuminated charge-coupled device (DBI-CCD) detector. The instrument was set at 1300W for Ca (RF power) 15 L/min plasma flow rate, 0.2 L/min Auxillary flow rate, 0.8 L/min nebuliser flow rate, radially viewed Ar-ICP. Source equilibration delay of 15 seconds. 2 mL/min sample and wash solution flow rates. Wash time 2 mL/min for 30 seconds between every sample and standard. For P ions, the instrument was set as above, except axially viewed Ar-ICP and 1400 W (RF power) and 0.4 L/min Nebuliser flow rate. Wavelengths used for Calcium were: 393.4 nm, 396.8 nm, 422.7 nm, wavelengths for phosphorus were 213.6 nm and 214.9 nm, wavelength for magnesium were: (II) 279.6 nm, (II) 280.3 nm, (I) 285.2 nm. Three independent replicates were tested for each wavelength and Scandium was used as internal standard at 1ppm monitored at 361 nm. The wash solution, sample and standard solutions were prepared with 2% nitric acid (prepared with Fisher Scientific S. G. 1.42 70% Laboratory Reagent Grade).

### Sound-based hydrogel membrane bio-assembly

The sound-based assembly of PA-**E3B**-Fibrin-CBP hydrogel composites started by pipetting 180 µL of PA-**E3B**-fibrinogen solution into the containers of squared or circular shape, depending on the geometry of the liquid receptacle. CBPs (0.5 mg) were added to the solution with a spatula and homogenously dispersed across the area of the container. After the CBPs were added, vibrations were applied by tuning the amplitude and frequencies until the patterned structures appeared, as described by Tognato *et al*. After formation of stable CBP patterns within the PA-**E3B**-fibrinogen solutions, 30µL of thrombin was added dropwise to allow slow gelation of the PA-**E3B**-fibrin-CBP composites and these were left undisturbed at room temperature for 30 mins before handling.

### Numerical simulations of the liquid surface displacement by Faraday waves

Numerical simulations of PA-**E3B**-fibrinogen surface displacement driven by hydrodynamic drag forces, were based on the fluid dynamics framework described by Ren *et al.* and described previously^[57]^. In brief, this framework reveals that CBPs under sound waves tend to accumulate in regions of minimal potential energy driven by the formation of Faraday waves, which produce standing wave patterns characterized by alternating nodes (areas of minimal displacement) and antinodes (areas of maximal displacement). Displacement at the liquid-air interface were mathematically modelled using Bessel functions of the first kind^[58,59]^. All simulations were conducted using Python 3.9 with the NumPy and Matplotlib libraries on a personal computer equipped with an 11th Gen Intel(R) i5-1145G7 processor and 16 GB of RAM.

### Mechanical tensile tests

Tensile tests of hydrogels were performed on an Instron 5969 testing machine, loading cell of 100N. Samples were positioned carefully between the clamps and tested at strain rate of 0.5 mm/min at room temperature. The hydrogels’ shape was approximated to a parallelepiped with area of 20 x 20 mm^2^ and thickness of 4 mm. Young’s moduli were extracted from the linear elastic region of force vs displacement curves and reported as average ± standard deviation of three independent samples.

### Finite Element Modelling (FEM)

The geometries for the FEM simulations were derived from experimental images of membranes patterned under acoustic stimulation at the following frequencies: 25, 57, 101, and 143 Hz. These images were binarized using ImageJ image processing software, and the 2D distribution of the PA phase within the fibrin hydrogel matrix was extracted by identifying relevant points. Constrained spline curves were generated by connecting these points using the Sketcher toolbox in the finite element software ABAQUS. The resulting geometries were then meshed using linear triangular finite elements and used for mechanical simulations. Each model consisted of two material phases: a) A soft matrix phase representing the fibrin hydrogel, and b) A stiffer inclusion phase representing the PA agglomerates. The 0 Hz condition was also simulated by randomly assigning approximately 50% of the elements the CBP properties and the remaining 50% the PA-fibrin properties. Due to the irregularity of the mesh, i.e., triangular elements of different areas, this assignment resulted in 53% of the 2D mesh area being modeled as CBP phase and the remaining 47% as fibrin phase. These proportions were considered reasonable as it should be noted that the experimental area covered by CBP at 0Hz cannot be accurately assessed due to the random placement of these particles during mixing. All models were square, like the experimental membranes, with an edge of 20 mm. The experimental accumulation of PA particles in the outer perimeter of the membrane was also simulated by adding a 0.3 mm thin frame. In constructing these models, care was taken to reproduce the experimental pattern area, i.e., the surface area fraction of the two phases. The models were subjected to uniaxial tensile loading by applying a uniform tensile force along one edge while constraining the opposite edge. Planar stress conditions were assumed. The FE analysis was static (i.e., no inertia effect) and accounted for geometric nonlinearities during the analysis step. The material properties were linear elastic. The Young’s modulus of the CBP phase was assigned to ensure numerical convergence under the simulated tensile load. Since it is experimentally difficult to measure the Young’s modulus of each phase, the modulus of the fibrin hydrogel was treated as a tunable parameter by setting the ratio 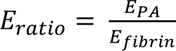. A sensitivity analysis was performed over a range of E_ration_ values to investigate their effect on the effective mechanical response of the patterned membranes.

### Polarized light microscopy

Polarised light microscopy was used to image alignments induced in the composite gels after application of sound waves. Sound-patterned hydrogels were fabricated as described above and imaged with a Zeiss Axioplan microscope equipped with a cross-polarizer.

### Fluorescence light microscopy of CBPs

To image the arrangement of CBPs within sound-assembled hydrogels, 10 mg of CBPs were dispersed into 1mL of PBS and incubated with 1 µL of Calcein AM (4µM, Invitrogen L3224) for 2 h at 4°C under constant agitation. After incubation, CBPs were retrieved from the solution with a Falcon cell strainer of 40µm mesh size and left to air dry at room temperature overnight. Calcein-stained CBPs were then dispersed into composite hydrogels as described above and imaged with a Leica SP8 upright dipping lens confocal microscope with excitation filter of 495 nm (green, Alexa Flour).

### Human Mesenchymal Stromal Cell (hMSC) culture

Patient-derived hMSCs were previously isolated and expanded from vertebral bone marrow samples (Bern Req-2023-00198) as previously described. Cells were cultured and further expanded in high-glucose Dulbecco’s-modified Eagle medium (Gibco, ThermoFisher, Zürich, Switzerland) with the addition of fetal bovine serum (Corning, Corning, NY, USA) and penicillin, streptomycin (P/S) (Gibco), with a final concentration of 10% and 1%, respectively. Once the cells reached 90% confluence, the medium was removed, and PBS was added to wash out completely the remaining medium.

Subsequently, PBS was removed, and 0.05% Trypsin/EDTA (Gibco) was added. Cells were then incubated at 37 °C for 5 min. After the incubation period, complete medium was added to inactivate Trypsin’s action. Detached cells re-suspended in the medium were then centrifuged at 700 rpm for 5 min. Subsequently, the supernatant was discharged, and cells were re-suspended in 1 mL of complete medium to proceed with cell counting. For MSC spheroids, cells were suspended in high-glucose Dulbecco’s-modified Eagle medium and cultured on a U-bottom 96-well culture plate.

### hMSC migration and infiltration study

To assess the cell migration and infiltration of hMSCs through the fabricated PA-fibrin-CBP hydrogels, hMSCs were stained with PKH cell membrane staining (code: PKH26GL, Merck, UK) and seeded on the hydrogels with a final concentration of 1×10^5^ cells/mL for 48 h. At each time point (4, 24, 48 h), cells were imaged with a Zeiss LSM800 confocal microscope (λ_ex_ 551 nm; λ_em_ 567 nm, PKH26 dye).

### hMSC viability and metabolic activity

Cell viability of MSCs seeded on the composite hydrogels was assessed using a standard live/dead staining protocol. After carefully removing the supernatant, cells were rinsed twice with PBS. A staining solution containing Calcein-AM (green; 2 × 10⁻³ M) and ethidium homodimer-1 (red; 4 × 10⁻³ M), both from Sigma-Aldrich, was then applied. Samples were incubated in the dark at 37 °C for 30 minutes. Post-incubation, the staining solution was removed, and samples were washed again with PBS. Fresh complete culture medium was added, and imaging was carried out using an EVOS M7000 fluorescence microscope (ThermoFisher Scientific, Waltham, USA). Metabolic activity of MSCs seeded on composite hydrogels was assessed with PrestoBlue® assay (code: A13261, ThermoFisher Scientific). Composite gels were formed in 96-well plates, MSCs were seeded on hydrogels and incubated with a mixture containing 90 µL of cell culture media plus 10 µL of PrestoBlue® reagent for 10 mins at 37 °C and 5% CO_2_. Following incubation, media was collected from the wells and fluorescence was measured using a Tecan Infinite® M-Plex microplate reader (excitaton wavelength: 560 nm; emission wavelength: 590 nm).

### Radial Intensity Profile and Colocalization Analysis

To evaluate the radial distribution and colocalization of fluorescence signals within patterned regions, we employed a radial intensity profile approach using the Radial Profile plugin in Fiji (ImageJ). Z-stack images were first projected into a single composite 2D image via maximum intensity projection, yielding a representative image for each time point. A circular region of interest (ROI) was then manually drawn around one of the four pattern nodes (circular features), based on visible signal distribution. The same ROI dimensions were subsequently applied to the remaining three pattern nodes within the same sample to ensure consistent sampling across all four regions. Radial intensity profiles were extracted for each ROI, and intensity values were normalized relative to a reference intensity point, typically corresponding to the local maximum or minimum within each pattern. This normalization step allowed comparison of spatial intensity distributions across multiple samples and time points. All results are reported as mean ± standard deviation, calculated from the four ROIs per sample (n = 4 per time point). Data analysis and plotting were performed using GraphPad Prism.

### Peripheral Blood Mononuclear Cell (PBMC) Isolation and Culture

Peripheral blood mononuclear cells (PBMCs) were isolated using a density gradient centrifugation method. Briefly, blood samples collected in lithium-heparin tubes were diluted with PBS (1:1 for whole blood) and gently layered onto Biocoll at room temperature. After centrifugation at 800 g for 20 minutes at room temperature with reduced acceleration and brake settings, the mononuclear cell layer at the interphase was carefully collected. The isolated cells were washed twice with cold PBS containing 2 mM EDTA, followed by centrifugation at 780 g and 300 g, respectively, at 8°C. Finally, the PBMC pellet was resuspended in cold complete RPMI medium, passed through a 70 µm cell strainer, and counted using a Neubauer chamber. For culture of PBMCs, 100 μL of PA-**E3**-Fibrin and PA-**E3B**-fibrin hydrogels were prepared into a 96-well plate format, as described in Section 5.2, and the well plate was incubated at 37°C for 30 minutes and left undisturbed to permit complete gel formation. Once gel formation was complete, PBMCs suspended in RPMI 1640 medium were added onto the surface of the gels within the 96-well plates and left to adhere.

### Targeted proteomics and bioinformatics analysis

PBMCs were cultured on gels made of fibrin or PA-**E3**-fibrin and PA-**E3B**-fibrin for evaluating targeted proteomic alterations induced by the cell-hydrogel interactions. In details, PBMC cultures, containing 200,000 cells per well in 96-well plates, were exposed to the hydrogels and incubated for 48 hours at 37°C in a CO₂ incubator. Following incubation, samples were collected for targeted proteomics using the Olink Proximity Extension Assay to explore immune and inflammatory responses (Olink Target 96, Inflammation and Immune Response panels, Olink Bioscience, Uppsala, Sweden). Proteomic profiles generated using Olink NPX technology were analyzed using R (v4.4.2). Data import and initial processing were performed using the OlinkAnalyze package (v4.2.0). After removing measurements flagged for low quality and merging the data with experimental metadata, rigorous quality control was conducted. This included assessing data distributions, identifying outliers, and validating the consistency of NPX scaling. To identify differentially expressed proteins, statistical models appropriate for the experimental design were selected; specifically, Welch’s two-sample t-tests were applied for comparisons between independent groups. Multiple comparisons were addressed by controlling the false discovery rate (FDR) using the Benjamini–Hochberg procedure, with significance defined as FDR < 0.05. Key differential expression results were visualized using volcano plots generated with ggplot2, highlighting effect sizes and statistical significance, and through heatmaps illustrating protein expression patterns across samples.

## Supporting Information

Supporting Information is available from the Wiley Online Library or from the author.

## Acknowledgements

The work was financially supported by AO Foundation, AO CMF (project: AOCMF-21-04S). AO CMF is a clinical division of the AO Foundation — an independent medically guided not-for-profit organization. AM acknowledges the Medical Research Council (UK Regenerative Medicine Platform Hub Acellular Smart Materials 3D Architecture, MR/R015651/1), the ERC PoC Grant NOVACHIP, and the NIHR Nottingham Biomedical Research Centre at University of Nottingham, Nottingham, UK.

## Data Availability Statement

The data that support the findings of this study are available from the corresponding author upon reasonable request.

## Supplementary Information

**Figure S1.**
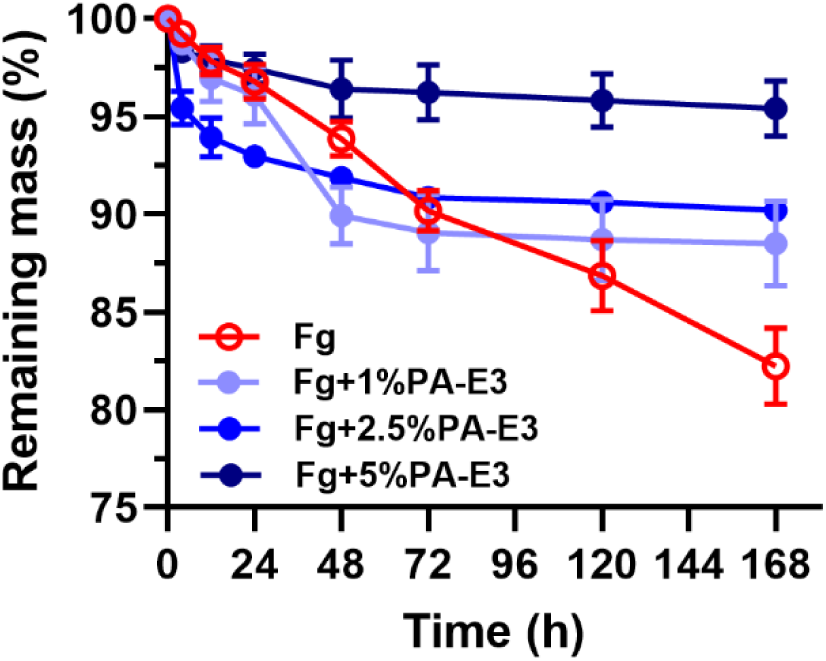
Enzymatic hydrogel degradation. Degradation over time of fibrin (Fg) and hybrid fibrin gels obtained by mixing fibrinogen with increasing concentrations of PA-E3 before addition of thrombin.

**Figure S2.**
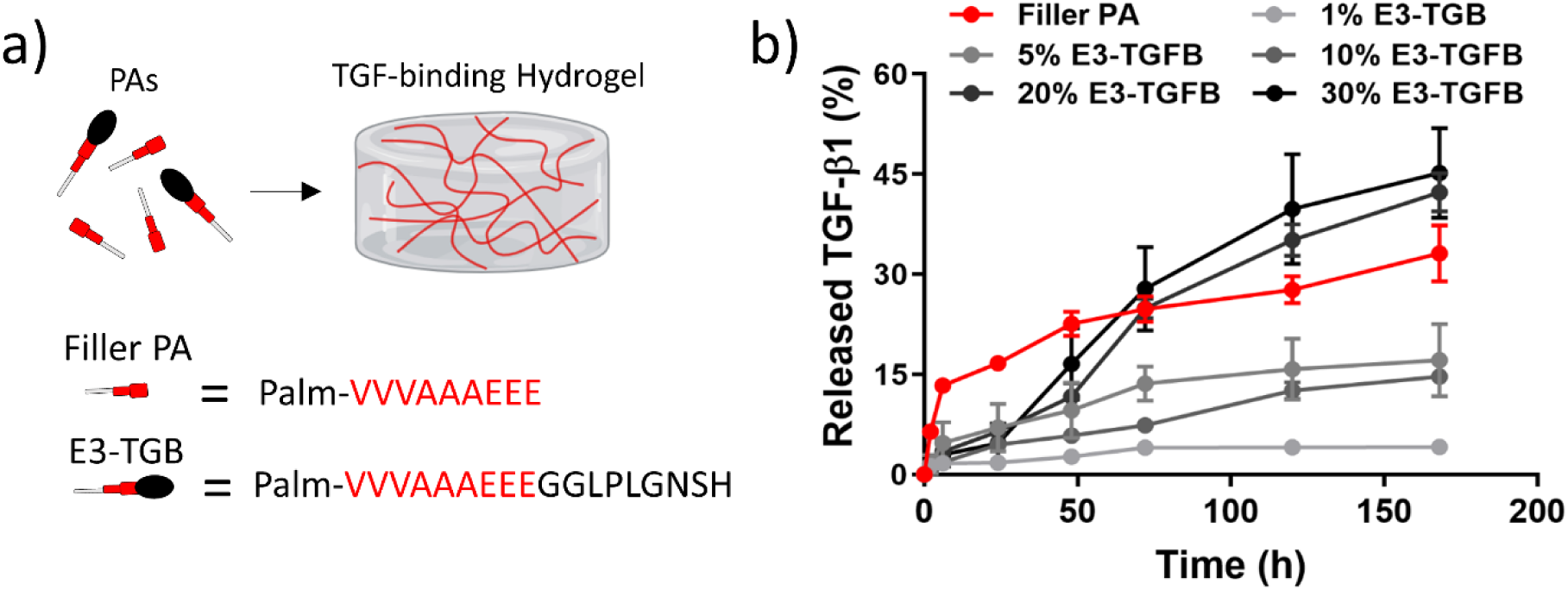
Release of bound TGFβ1. a) PA sequences used to bind and release TGFβ1. b) ELISA assays of TGFβ1 released from 3D hydrogels containing PA-E3 only (‘filler PA’) and gels containing increasing weight ratios of TGFβ1-binding peptides (‘E3-TGB’, i.e. PA-E3 displaying the GF-binding peptide sequence ‘LPLGNSH’). Graphs show dependence of release profiles on epitope density.

**Figure S3.**
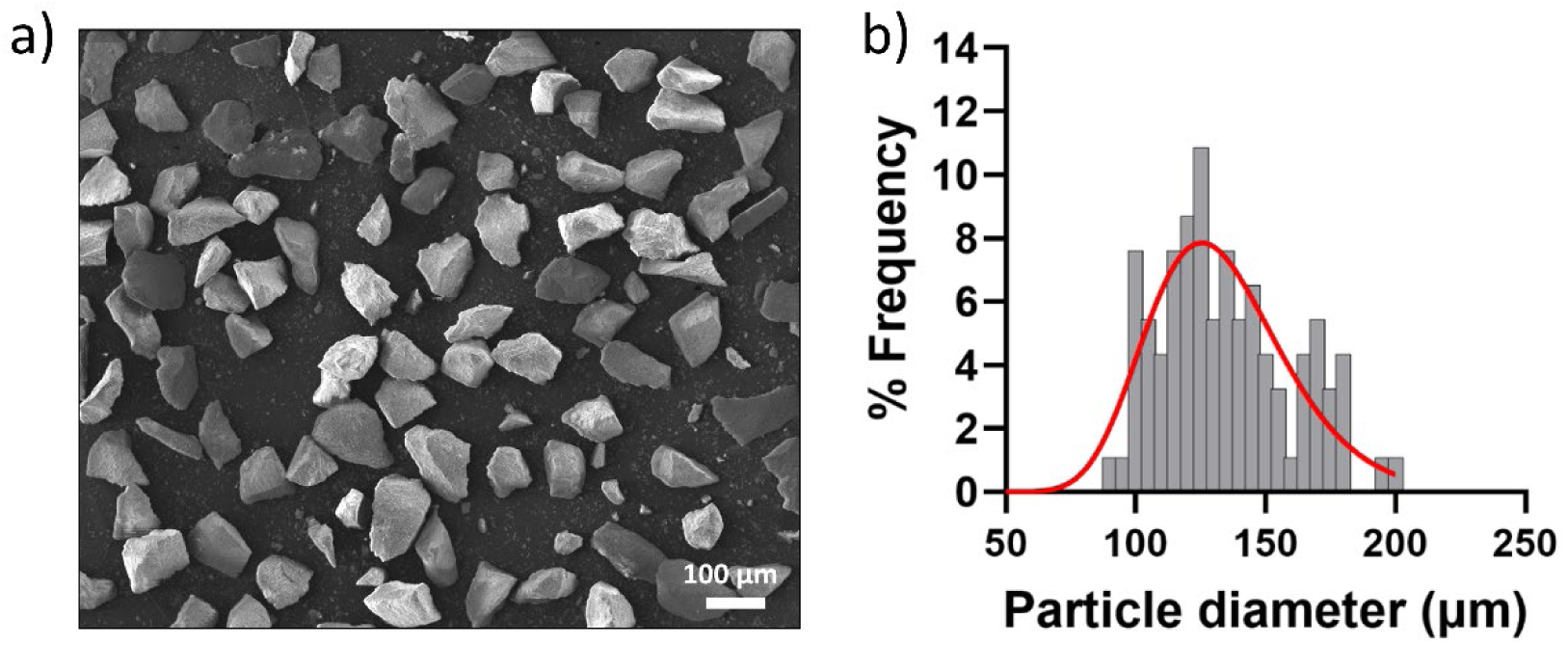
Size distribution of calcined bone particles (CBPs). a) SEM image of CBPs on a carbon tape. b) Frequency distribution of the CBP size shows the majority of particles lying between 80 and 180 µm in size.

**Figure S4.**
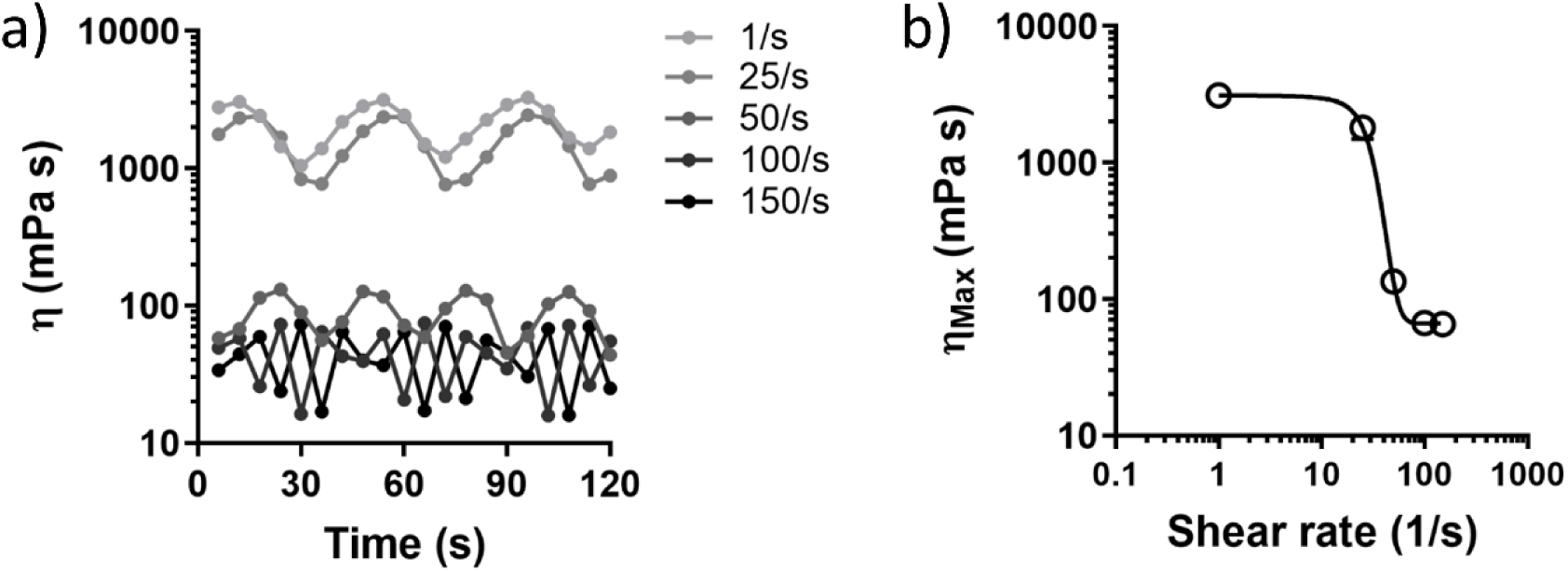
Rheological behavior of PA-fibrinogen solution. a) Flow sweep measurements of PA-fibrinogen solution recording viscosity () while varying time and frequency of test oscillations. b) Maximum viscosity extracted from the maxima of graph a) in function of shear rates. Both graphs shows time-dependent and shear-thinning behaviour of the PA-fibrinogen solution used

